# *Listeria monocytogenes* requires cellular respiration for NAD^+^ regeneration and pathogenesis

**DOI:** 10.1101/2021.11.25.470024

**Authors:** Rafael Rivera-Lugo, David Deng, Andrea Anaya-Sanchez, Sara Tejedor-Sanz, Valeria M Reyes Ruiz, Hans B Smith, Denis V Titov, John-Demian Sauer, Eric P Skaar, Caroline M Ajo-Franklin, Daniel A Portnoy, Samuel H Light

## Abstract

Cellular respiration is essential for multiple bacterial pathogens and a validated antibiotic target. In addition to driving oxidative phosphorylation, bacterial respiration has a variety of ancillary functions that obscure its contribution to pathogenesis. We find here that the intracellular pathogen *Listeria monocytogenes* encodes two respiratory pathways which are partially functionally redundant and indispensable for pathogenesis. Loss of respiration decreased NAD^+^ regeneration, but this could be specifically reversed by heterologous expression of a water-forming NADH oxidase (NOX). NOX expression fully rescued intracellular growth defects and increased *L. monocytogenes* loads >1,000-fold in a mouse infection model. Consistent with NAD^+^ regeneration maintaining *L. monocytogenes* viability and enabling immune evasion, a respiration-deficient strain exhibited elevated bacteriolysis within the host cytosol and NOX rescued this phenotype. These studies show that NAD^+^ regeneration, rather than oxidative phosphorylation, represents the primary role of *L. monocytogenes* respiration and highlight the nuanced relationship between bacterial metabolism, physiology, and pathogenesis.

## Introduction

Distinct metabolic strategies allow microbes to extract energy from diverse surroundings and colonize nearly every part of the earth. Microbial energy metabolisms vary greatly but can be generally categorized as possessing fermentative or respiratory properties. Cellular respiration is classically described by a multi-step process that initiates with the enzymatic oxidation of organic matter and the accompanying reduction of NAD^+^ (nicotinamide adenine dinucleotide) to NADH. Respiration of fermentable sugars typically starts with glycolysis, which generates pyruvate and NADH. Pyruvate then enters the tricarboxylic acid (TCA) cycle, where its oxidation to carbon dioxide is coupled to the production of additional NADH. NADH generated by glycolysis and the TCA cycle is then oxidized by NADH dehydrogenase to regenerate NAD^+^ and the resulting electrons are transferred via an electron transport chain to a terminal electron acceptor. While mammals strictly rely upon aerobic respiration, which uses oxygen as the terminal electron acceptor, microbes reside in diverse oxygen-limited environments and have varying and diverse capabilities to use disparate non-oxygen respiratory electron acceptors. Whatever the electron acceptor, electron transfer in the electron transport chain is coupled to proton pumping across the bacterial inner membrane. This generates a proton gradient or proton motive force, which powers a variety of processes, including ATP production by ATP synthase.

Respiratory pathways are important for several aspects of bacterial physiology. Respiration’s role in establishing the proton motive force allows bacteria to generate ATP from non-fermentable energy sources (which are not amenable to ATP production by substrate-level phosphorylation) and increases ATP yields from fermentable energy sources. In addition to these roles in ATP production, the proton motive force is directly involved in many other aspects of bacterial physiology, including the regulation of cytosolic pH, transmembrane solute transport, ferredoxin-dependent metabolisms, protein secretion, and flagellar motility.^1–6^ Beyond the proton motive force, respiration functions to regenerate NAD^+^, which is essential for enabling the continued function of glycolysis and other metabolic processes. By obviating fermentative mechanisms of NAD^+^ regeneration, respiration increases metabolic flexibility, which, among other metabolic consequences, can enhance ATP production by substrate-level phosphorylation.^7^

Bacterial pathogens reside within a host where they must employ fermentative or respiratory metabolisms to power growth. Pathogen respiratory processes have been linked to host-pathogen conflict in several contexts. Phagocytic cells target bacteria by producing reactive nitrogen species that inhibit aerobic respiration.^8^ *Aggregatibacter actinomycetemcomitans*, *Salmonella enterica*, *Streptococcus agalactiae*, and *Staphylococcus aureus* mutants with impaired aerobic respiration are attenuated in murine models of systemic disease.^9–12^ Aerobic respiration is vital for *Mycobacterium tuberculosis* pathogenesis and persister cell survival, making respiratory systems validated anti-tuberculosis drug targets.^13,14^ Respiratory processes that use oxygen, tetrathionate, and nitrate as electron acceptors are important for the growth of *Salmonella enterica* and *Escherichia coli* in the mammalian intestinal lumen.^15–17^ While several studies have linked respiration in bacterial pathogens to the use of specific electron donors (i.e., non-fermentable energy sources) within the intestinal lumen, the particular respiratory functions important for systemic bacterial infections remain largely unexplained.^18–22^

*Listeria monocytogenes* is a human pathogen that, after being ingested on contaminated food, can gain access to the host cell cytosol and use actin-based motility to spread from cell-to-cell.^23^ *L. monocytogenes* has two respiratory-like electron transport chains. One electron transport chain is dedicated to aerobic respiration and uses QoxAB (*aa*_3_) or CydAB (*bd*) cytochrome oxidases for terminal electron transfer to O_2_ (**Fig 1a**).^24^ We recently identified a second flavin-based electron transport chain that transfers electrons to extracytosolic acceptors (including ferric iron and fumarate) and can promote growth in anaerobic conditions (**Fig 1a**).^25–27^

**Figure 1.**
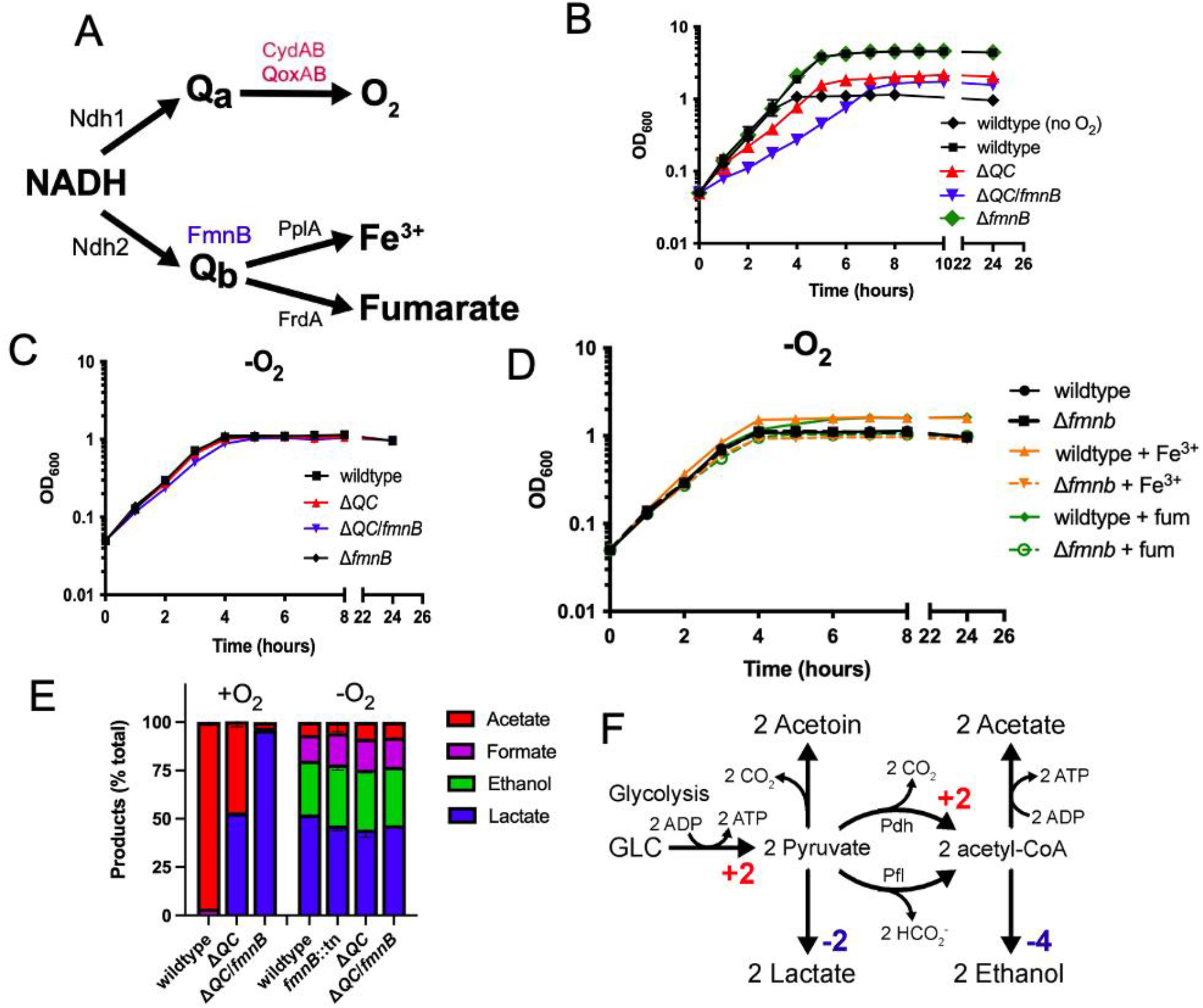
Respiration impacts *L. monocytogenes* growth and fermentative output. (A) Proposed respiratory electron transport chains in *L. monocytogenes*. Different NADH dehydrogenases likely transfer electrons to distinct but presently unidentified quinones (Q_a_ and Q_b_). FmnB catalyzes assembly of essential components of the electron transport chain, PplA and FrdA, that can transfer electrons to ferric iron and fumarate, respectively. Other proteins involved in the terminal electron transfer steps are noted. (B) Optical density of *L. monocytogenes* strains aerobically grown in nutrient-rich media, with the anaerobically grown wildtype strain provided for context. The means and standard deviations from three independent experiments are shown. (C) Optical density of *L. monocytogenes* strains grown anaerobically in nutrient-rich media. The data represent the means and standard deviations from three independent experiments. (D) Optical density of anaerobically grown strains in nutrient-rich media supplemented with the alternative electron acceptors ferric iron (Fe^3+^) or fumarate (fum), as indicated. The means and standard deviations from three independent experiments are shown. (E) Fermentation products of *L. monocytogenes* strains grown in nutrient-rich media under aerobic and anaerobic conditions. Error bars show standard deviations. Results from three independent experiments are shown. (F) Proposed pathways for *L. monocytogenes* sugar metabolism. The predicted number of NADH generated (+) or consumed (-) in each step is indicated. PplA, peptide pheromone-encoding lipoprotein A; FrdA, fumarate reductase A; Δ*QC*, Δ*qoxA*/Δ*cydAB*; Δ*QC*/*fmnB*, Δ*qoxA*/Δ*cydAB*/*fmnB*::tn; GLC, glucose; Pdh, pyruvate dehydrogenase; Pfl, pyruvate formate-lyase.

Despite lacking a complete TCA cycle, previous studies have shown that aerobic respiration is important for systemic spread of *L. monocytogenes*.^24,28–30^ Saccharolytic microbes that similarly contain a respiratory electron transport chain but lack a complete TCA cycle can be considered to employ a respiro-fermentative metabolism.^31^ Respiro-fermentative metabolisms tune the cell’s fermentative output and often manifest with the respiratory regeneration of NAD^+^ enabling a shift from the production of reduced (e.g., lactic acid and ethanol) to oxidized (e.g., acetic acid) fermentation products. In respiro-fermentative lactic acid bacteria that are closely related to *L. monocytogenes*, cellular respiration results in a modest growth enhancement, but is generally dispensable. The role of aerobic respiration for *L. monocytogenes* pathogenesis thus might be considered surprising and remains unclear.^31,32^

The studies presented here sought to address the role of respiration in *L. monocytogenes* pathogenesis. Our results confirm tha*t L. monocytogenes* exhibits a respiro-fermentative metabolism and show that its two respiratory systems are partially functionally redundant under aerobic conditions. We find that the respiration-deficient *L. monocytogenes* strains exhibit severely attenuated virulence and lyse within the cytosol of infected cells. Finally, we selectively abrogate the effect of diminished NAD+ regeneration in respiration-deficient *L. monocytogenes* strains by heterologous expression of a water-forming NADH oxidase (NOX) and find that this restores virulence. These results thus elucidate the basis of *L. monocytogenes* cellular respiration and demonstrate that NAD^+^ regeneration represents a key function of this activity in *L. monocytogenes* pathogenesis.

## Results

### *L. monocytogenes*’ electron transport chains have distinct roles in aerobic and anaerobic growth

*We* selected previously characterized Δ*qoxA*/Δ*cydAB* and *fmnB*::tn *L. monocytogenes* strains to study the role of aerobic respiration and extracellular electron transfer, respectively.^25,30^ In addition, we generated a *ΔqoxA/ΔcydAB/fmnB*::tn *L. monocytogenes* strain to test for functional redundancies of aerobic respiration and extracellular electron transfer. Initial studies measured the growth of these strains on nutritionally rich media (with glucose as the primary growth substrate) in the presence/absence of electron acceptors.

Compared to anaerobic conditions that lacked an electron acceptor, we found that aeration led to a relatively modest increase in growth of wildtype and *fmnB*::tn strains (**Fig.1b & c**). This growth enhancement could be attributed to aerobic respiration, as aerobic growth of the Δ*qoxA*/Δ*cydAB* strain resembled anaerobically cultured strains (**Fig.1b & c**). Similarly, in anaerobic conditions, inclusion of the extracellular electron acceptor ferric iron resulted in a small growth enhancement of wildtype *L. monocytogenes* (**Fig.1d**). This phenotype could be attributed to extracellular electron transfer, as ferric iron failed to stimulate growth of the *fmnB*::tn strain (**Fig 1d**). These findings are consistent with aerobic respiration and extracellular electron transfer possessing distinct roles in aerobic and anaerobic environments, respectively.

The Δ*qoxA*/Δ*cydAB*/*fmnB*::tn strain exhibited the most striking growth pattern, since it lacked a phenotype under anaerobic conditions but had impaired aerobic growth, even relative to the Δ*qoxA*/Δ*cydAB* strain (**Fig 1b & 1c)**. Notably, Δ*qoxA*/Δ*cydAB*/*fmnB*::tn was the sole strain tested with a substantially reduced growth rate in the presence of oxygen (**Fig 1b & 1c**). These observations suggest that aerobic extracellular electron transfer activity can partially compensate for the loss of aerobic respiration and that oxygen inhibits *L. monocytogenes* growth in the absence of both electron transport chains.

### Respiration alters *L. monocytogenes*’ fermentative output

Respiration is classically defined by the use of the TCA cycle to fully oxidize an electron donor (e.g., glucose) to carbon dioxide. However, *L. monocytogenes* lacks a TCA cycle and instead converts sugars into multiple fermentation products.^33^ We thus asked how respiration impacts *L. monocytogenes*’ fermentative output. Under anaerobic conditions that lacked an alternative electron acceptor, *L. monocytogenes* exhibited a pattern of mixed acid fermentation, with lactic acid being most abundant and ethanol, formic acid, and acetic acid being produced at lower levels (**Fig 1e**). By contrast, under aerobic conditions *L. monocytogenes* almost exclusively produced acetic acid (**Fig 1e**). Consistent with respiration being partially responsible for the distinct aerobic vs. anaerobic responses, Δ*qoxA*/Δ*cydAB* and Δ*qoxA/ΔcydAB/fmnB::*tn strains failed to undergo drastic shifts in fermentative output when grown in aerobic conditions. The Δ*qoxA*/Δ*cydAB* strain mainly produced lactic acid in the presence of oxygen and this trend was even more pronounced in the Δ*qoxA*/Δ*cydAB*/*fmnB*::tn strain, which almost exclusively produced lactic acid (**Fig 1e**). These results show that aerobic respiration induces a shift to acetic acid production and support the conclusion that *L. monocytogenes*’ two electron transport chains are partially functionally redundant in aerobic conditions.

A comparison of fermentative outputs across the experimental conditions also clarifies the basis of central energy metabolism in *L. monocytogenes*. A classical glycolytic metabolism in *L. monocytogenes* likely generates ATP and NADH. In the absence of oxygen or an alternative electron acceptor, NAD^+^ is regenerated by coupling NADH oxidation to the reduction of pyruvate to lactate or ethanol. In the presence of oxygen, NADH oxidation is coupled to the reduction of oxygen and pyruvate is converted to acetate. Moreover, the pattern of anaerobic formate production is consistent with aerobic acetyl-CoA production through pyruvate dehydrogenase and anaerobic production through pyruvate formate-lyase (**Fig 1f**). Collectively, these observations suggest that *L. monocytogenes* prioritizes balancing NAD^+^/NADH levels in the absence of an electron acceptor and maximizing ATP production in the presence of oxygen. In the absence of oxygen, NAD^+^/NADH redox homeostasis is achieved by minimizing NADH produced in acetyl-CoA biosynthesis and by consuming NADH in lactate/ethanol fermentation (**Fig 1f**). In the presence of oxygen, ATP yields are maximized through respiration and increased acetate kinase activity (**Fig 1f**).

### Respiratory capabilities are essential for *L. monocytogenes* pathogenesis

We next asked about the role of cellular respiration in intracellular *L. monocytogenes* growth and pathogenesis. The *fmnB*::tn mutant deficient for extracellular electron transfer was previously shown to resemble the wildtype *L. monocytogenes* strain in a murine model of infection.^25^ We found that this mutant also did not differ from wildtype *L. monocytogenes* in growth in bone marrow-derived macrophages and a plaque assay that monitors bacterial growth and cell-to-cell spread (**Fig 2a & 2b**). Consistent with previous reports, the Δ*qoxA*/Δ*cydAB* strain deficient for aerobic respiration was attenuated in the plaque assay and murine model of infection, but resembled wildtype *L. monocytogenes* in macrophage growth (**Fig 2a–2c**).^24,30^ Combining mutations that resulted in the loss of both extracellular electron transfer and aerobic respiration produced even more pronounced phenotypes. The Δ*qoxA*/Δ*cydAB*/*fmnB*::tn strain did not grow intracellularly in macrophages and fell below the limit of detection in the plaque assay and murine infection model (**Fig 2a–2c**). These results thus demonstrate that respiratory activities are essential for *L. monocytogenes* virulence and that the organism’s two respiratory pathways are partially functionally redundant within a mammalian host.

**Figure 2.**
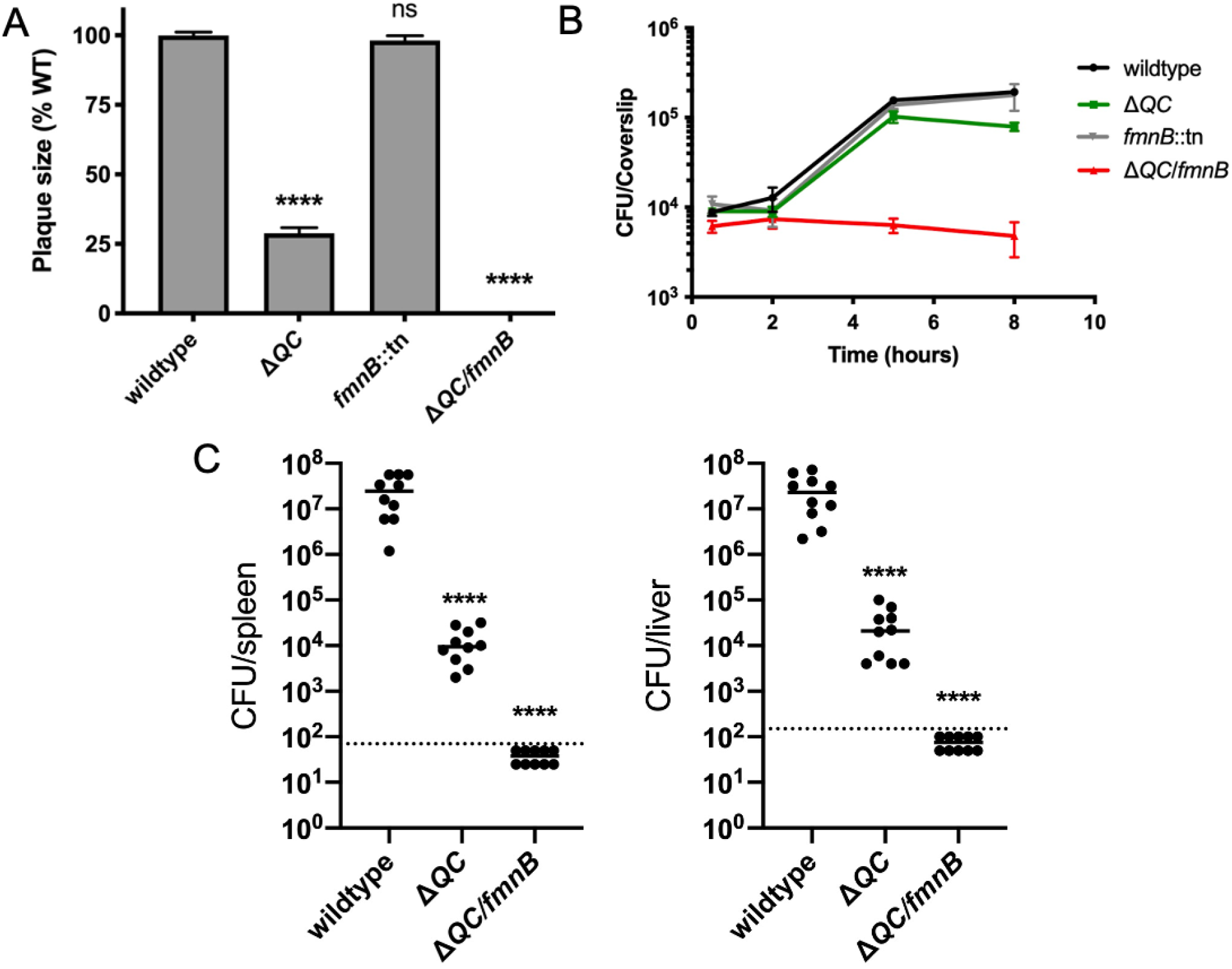
Respiration is required for *L. monocytogenes* virulence. (A) Plaque formation by cell-to-cell spread of *L. monocytogenes* strains in monolayers of mouse L2 fibroblast cells. The mean plaque size of each strain is shown as a percentage relative to the wildtype plaque size. Error bars represent standard deviations of the mean plaque size from two independent experiments. Statistical analysis was performed using one-way ANOVA and Dunnett’s post-test comparing wildtype to all the other strains. ****, *P* < 0.0001; ns, no significant difference (*P* > 0.05). (B) Intracellular growth of *L. monocytogenes* strains in murine bone marrow-derived macrophages (BMMs). One hour post-infection, infected BMMs were treated with 50 μg/mL of gentamicin to kill extracellular bacteria. Colony forming units (CFU) were enumerated at the indicated times. Results are representative of two independent experiments. (C) Bacterial burdens in murine spleens and livers 48 hours post-intravenous infection with indicated *L. monocytogenes* strains. The median values of the CFUs are denoted by black bars. The dashed lines represent the limit of detection. Data were combined from two independent experiments, *n* = 10 mice per strain. Statistical significance was evaluated using one-way ANOVA and Dunnett’s post-test using wildtype as the control. ****, *P* < 0.0001. Δ*QC*, Δ*qoxA*/Δ*cydAB*; Δ*QC/fmnB*, Δ*qoxA*/Δ*cydAB*/*fmnB*::tn.

### Expression of NOX restores NAD^+^ levels in *L. monocytogenes* respiration mutants

Cellular respiration both regenerates NAD^+^ and establishes a proton motive force that is important for various aspects of bacterial physiology. The involvement of respiration in these two distinct processes can confound the analysis of respiration-impaired phenotypes. However, the heterologous expression of water-forming NADH oxidase (NOX) has been used to decouple these functionalities in mammalian cells (**Fig 3a**).^34^ Because NOX regenerates NAD^+^ without pumping protons across the membrane, its introduction to a respiration-deficient cell can correct an NAD^+^/NADH imbalance, thereby isolating the role of the proton motive force in the phenotype.^34,35^

**Figure 3.**
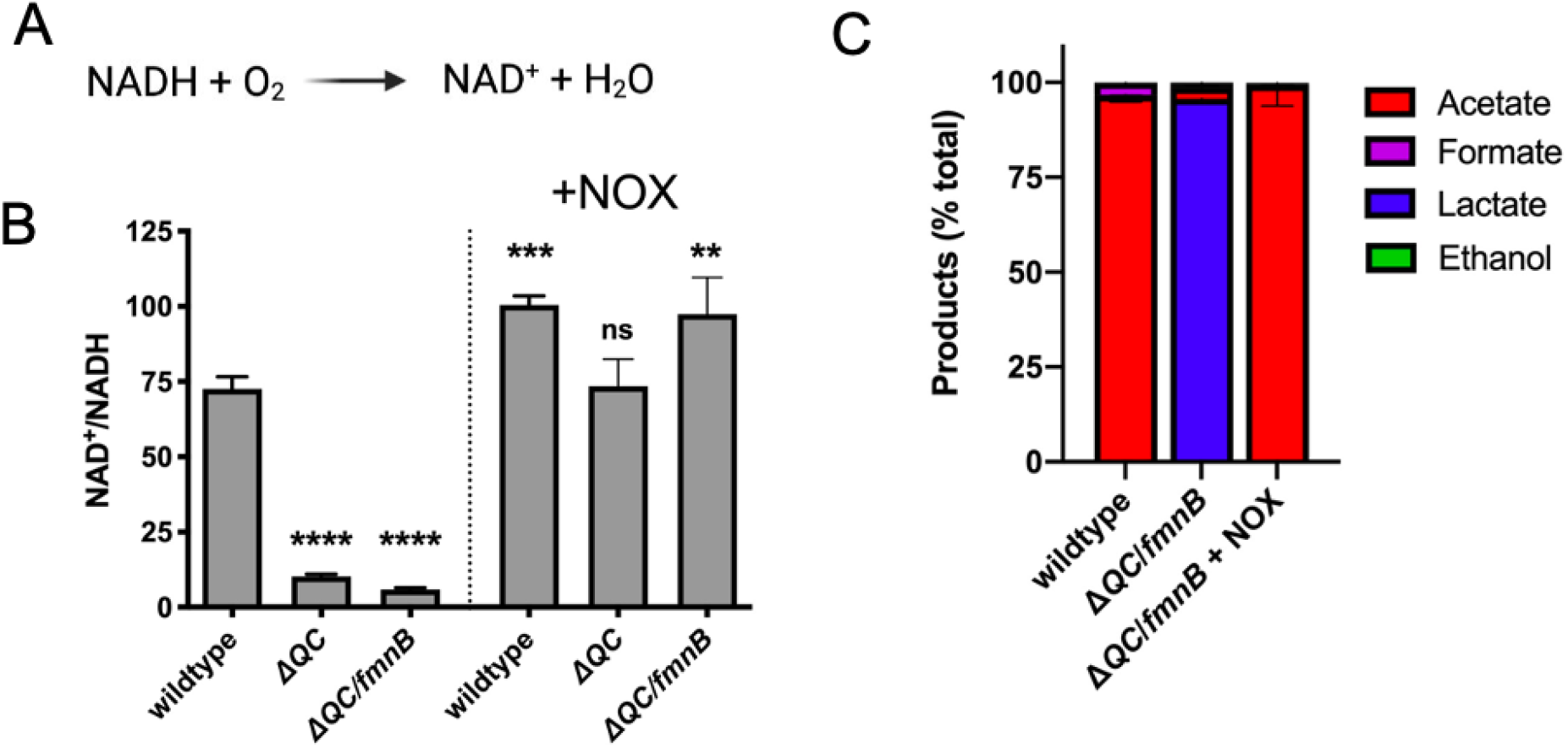
Water-forming NADH oxidase (NOX) restores redox homeostasis in respiration-deficient *L. monocytogenes* strains. (A) Reaction catalyzed by the *Lactococcus lactis* water-forming NOX, which is the same as aerobic respiration without the generation of a proton motive force. (B) NAD^+^/NADH ratios of parent and NOX-complemented *L. monocytogenes* strains grown aerobically in nutrient-rich media to mid-logarithmic phase. Results from three independent experiments are presented as means and standard deviations. Statistical significance was calculated using one-way ANOVA and Dunnett’s post-test using the wildtype parent strain as the control. ****, *P* < 0.0001; ***, *P* < 0.001; **, *P* < 0.01; ns, not statistically significant (*P* > 0.05). (C) Fermentation products of *L. monocytogenes* strains grown in nutrientrich media under aerobic conditions. Error bars show standard deviations. Results from three independent experiments are shown. Δ*QC*, Δ*qoxA*/Δ*cydAB*; Δ*QC/fmnB*, Δ*qoxA*/Δ*cydAB*/*fmnB*::tn; +NOX, strains complemented with *L. lactis* NOX.

To address which aspect of cellular respiration was important for *L. monocytogenes* pathogenesis, we introduced the previously characterized *Lactococcus lactis* water-forming NOX to the genome of respiration-deficient *L. monocytogenes* strains.^36–38^ We confirmed that the Δ*qoxA*/Δ*cydAB* and Δ*qoxA*/Δ*cydAB*/*fmnB*::tn strains exhibited decreased NAD^+^/NADH ratios and that constitutive expression of NOX rescued this phenotype (**Fig 3b**). We also found that NOX expression restored the predominance of acetic acid production to the aerobically grown Δ*qoxA*/Δ*cydAB*/*fmnB*::tn strain – confirming that the altered fermentative output of this respiration-deficient strain stems from impaired NAD^+^ regeneration (**Fig 3c**). These experiments demonstrate that NOX can be used as a tool to manipulate the NAD^+^/NADH ratio in bacteria.

### Respiration is critical for regenerating NAD^+^ during *L. monocytogenes* pathogenesis

We next sought to dissect the relative importance of respiration in generating a proton motive force versus maintaining redox homeostasis for *L. monocytogenes* virulence. We tested NOX-expressing Δ*qoxA*/Δ*cydAB* and Δ*qoxA*/Δ*cydAB*/*fmnB*::tn strains for macrophage growth, plaque formation, and in the murine infection model. Expression of NOX almost fully rescued the plaque assay and macrophage growth phenotypes of the Δ*qoxA*/Δ*cydAB* and Δ*qoxA*/Δ*cydAB*/*fmnB*::tn strains (**Fig 4a and 4b**). NOX expression also partially rescued *L. monocytogenes* virulence in the murine infection model (**Fig 4c**). Notably, NOX expression had a greater impact on the *L. monocytogenes* load in the spleen than the liver, suggesting distinct functions of respiration for *L. monocytogenes* colonization of these two organs (**Fig 4c**). These results are consistent with NAD^+^ regeneration representing the primary role of respiration in *L. monocytogenes* pathogenesis to an organ-specific extent.

**Figure 4.**
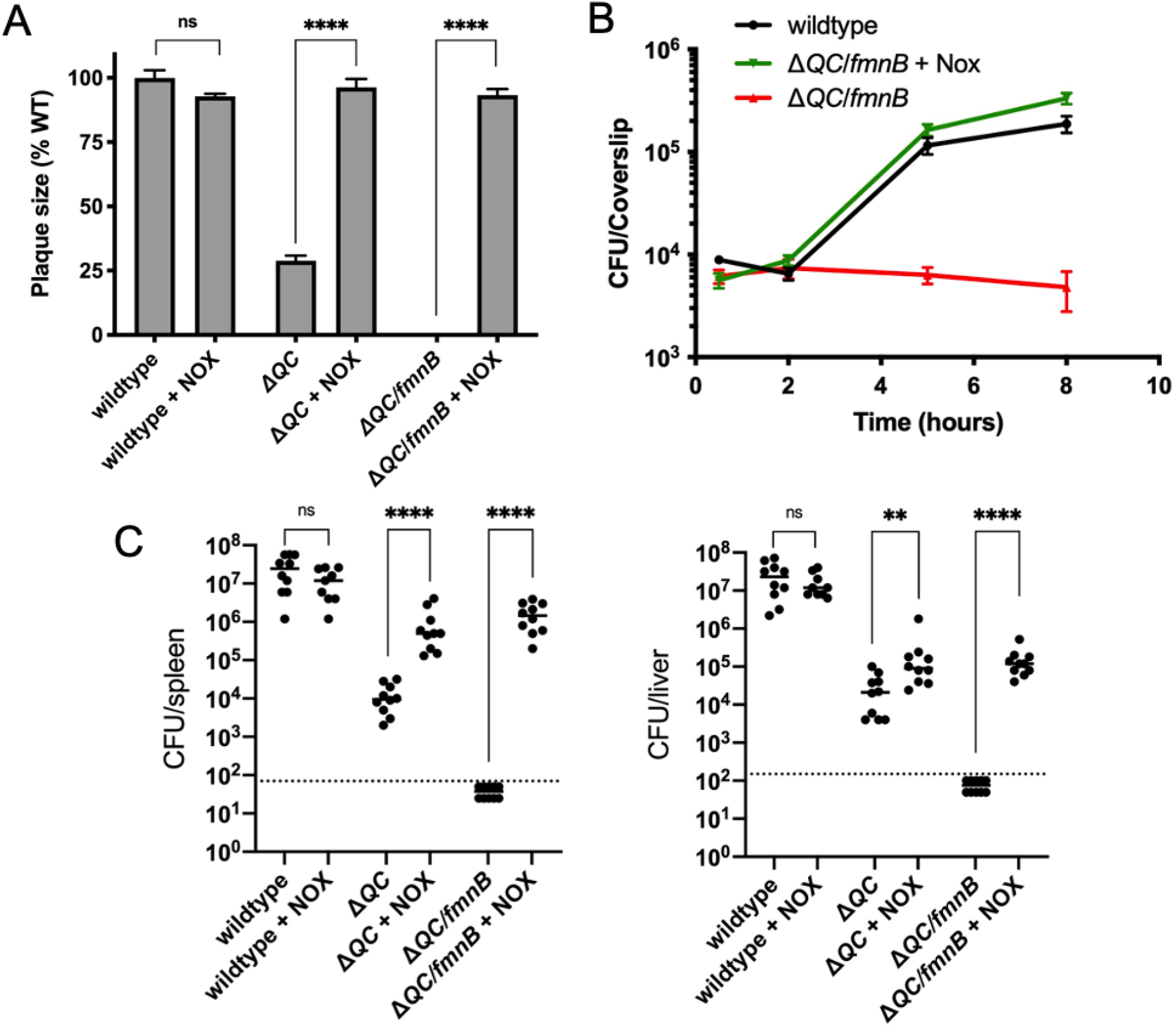
NOX expression restores virulence to respiration-deficient *L. monocytogenes* strains. (A) Plaque formation by cell-to-cell spread of *L. monocytogenes* strains in monolayers of mouse L2 fibroblast cells. The mean plaque size of each strain is shown as a percentage relative to the wildtype plaque size. Error bars represent standard deviations of the mean plaque size from two independent experiments. Statistical analysis was performed using the unpaired two-tailed *t* test. ****, *P* < 0.0001; ns, no significant difference (*P* > 0.05). (B) Intracellular growth of *L. monocytogenes* strains in murine bone marrow-derived macrophages (BMMs). One hour post-infection, infected BMMs were treated with 50 μg/mL of gentamicin to kill extracellular bacteria. Colony forming units (CFU) were enumerated at the indicated times. Results are representative of three independent experiments. (C) Bacterial burdens in murine spleens and livers 48 hours post-intravenous infection with indicated *L. monocytogenes* strains. The median values of the CFUs are denoted by black bars. The dashed lines represent the limit of detection. Data were combined from two independent experiments, *n*= 10 mice per strain, but for the wildtype + NOX strain (*n*= 9 mice). Statistical significance was evaluated using the unpaired two-tailed *t* test. ****, *P* < 0.0001; **, *P* < 0.01; ns, no significant difference (*P* > 0.05). Δ*QC*, Δ*qoxA*/Δ*cydAB*; Δ*QC*/*fmnB*, Δ*qoxA*/Δ*cydAB*/*fmnB*::tn; + NOX, strains complemented with *L. lactis* NOX.

### Impaired redox homeostasis is associated with increased cytosolic *L. monocytogenes* lysis

We next asked why respiration-mediated redox homeostasis was critical for *L. monocytogenes* pathogenesis. We reasoned previous descriptions of *L. monocytogenes* quinone biosynthesis mutants might provide a clue. Quinones are a family of redox-active cofactors that have essential functions in respiratory electron transport chains.^39^ Our previous studies suggested that distinct quinones function in flavin-based electron transfer and aerobic respiration^25^. A separate set of studies found that *L. monocytogenes* quinone biosynthesis mutants exhibited divergent phenotypes. *L. monocytogenes* strains defective in upstream steps of the quinone biosynthesis pathway exhibited increased bacteriolysis in the cytosol of host cells and were severely attenuated for virulence. By contrast, *L. monocytogenes* strains defective in downstream steps of the quinone biosynthesis pathway did not exhibit increased cytosolic bacteriolysis and had less severe virulence phenotypes^30,40,41^. These divergent phenotypic responses resemble the loss of aerobic respiration versus the loss of aerobic respiration *plus* flavin-based electron transfer observed in our studies. The distinct virulence phenotype of quinone biosynthesis mutants could thus be explained by the upstream portion of the quinone biosynthesis pathway being required for both aerobic respiration and flavin-based electron transfer, with the downstream portion of the pathway only being required for aerobic respiration (**Fig 5a**).

**Figure 5.**
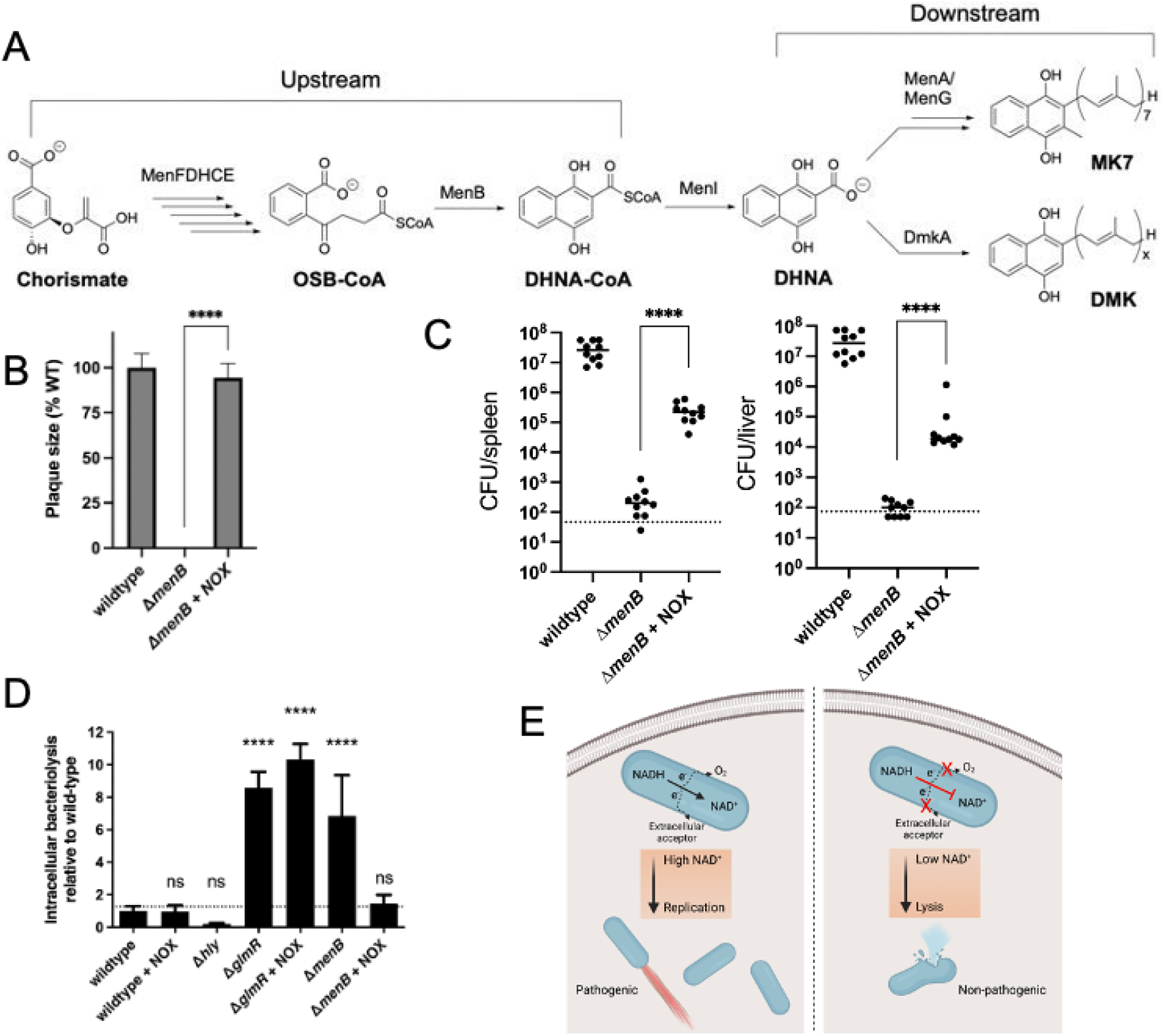
Impaired redox homeostasis accounts for elevated bacteriolysis of a respiration-deficient *L. monocytogenes* strain in the cytosol of infected cells. (A) Proposed *L. monocytogenes* quinone biosynthesis pathway. An unidentified demethylmenaquinone (DMK) is proposed to be required for the flavin-based electron transfer pathway and MK7 required for aerobic respiration. Loss of the upstream portion of the pathway is anticipated to impact both electron transport chains. (B) Plaque formation by cell-to-cell spread of *L. monocytogenes* strains in monolayers of mouse L2 fibroblast cells. The mean plaque size of each strain is shown as a percentage relative to the wildtype plaque size. Error bars represent standard deviations of the mean plaque size from two independent experiments. Statistical analysis was performed using the unpaired two-tailed *t* test. ****, *P* < 0.0001. (C) Bacterial burdens in murine spleens and livers 48 hours post-intravenous infection with indicated *L. monocytogenes* strains. The median values of the CFUs are denoted by black bars. The dashed lines represent the limit of detection. Data were combined from two independent experiments, *n*= 10 mice per strain. Statistical significance was evaluated using the unpaired two-tailed *t* test. ****, *P* < 0.0001. (D) Bacteriolysis of *L. monocytogenes* strains in bone marrow-derived macrophages. The data are normalized to wildtype bacteriolysis levels and presented as means and standard deviations from three independent experiments. Statistical significance was calculated using one-way ANOVA and Dunnett’s post-test using the wildtype parent strain as the control. ****, *P* < 0.0001; ns, no significant difference (*P* > 0.05). (E) Model of the role of respiration in *L. monocytogenes* pathogenesis. On the left, an intracellular bacterium with the ability to oxidize NADH and transfer electrons through the aerobic and EET electron transport chains can regenerate and maintain high NAD^+^ levels which allows the bacterium to grow and be virulent. On the right, an intracellular bacterium unable to regenerate NAD^+^, by lacking electron transport chains, is avirulent because it lyses in the cytosol of infected cells.

Based on the proposed roles of quinones in respiration, we hypothesized that the severe phenotypes previously described for the upstream quinone biosynthesis mutants were due to an imbalance in the NAD^+^/NADH ratio. To address this hypothesis, we first confirmed that the Δ*menB* strain, which is defective in upstream quinone biosynthesis, exhibited a phenotype similar to the Δ*qoxA*/Δ*cydAB*/*fmnB*::tn strain for plaque formation and in the murine infection model (**Fig 5b and 5c**). We next tested the effect of NOX expression on virulence phenotypes for the Δ*menB* strain. NOX expression rescued Δ*menB* phenotypes for plaque formation and in the murine infection model to a strikingly similar extent as the Δ*qoxA*/Δ*cydAB*/*fmnB*::tn strain (**Fig 5b and 5c**). These results thus provide evidence that quinone biosynthesis is essential for respiration and that the severity of the Δ*menB* phenotype is, in large part, to the role of respiration in regenerating NAD^+^.

Numerous adaptations allow *L. monocytogenes* to colonize the host cytosol, including a resistance to bacteriolysis. Minimizing bacteriolysis within the host cytosol is important to the pathogen because it can activate the host’s innate immune responses, including pyroptosis, a form of programmed cell death, which severely reduces *L. monocytogenes* virulence.^42^ *L. monocytogenes* strains deficient for the upstream quinone biosynthesis steps were previously identified as having an increased susceptibility to bacteriolysis in the macrophage cytosol.^30^ We thus hypothesized that decreased virulence of respiration-deficient strains might relate to increased cytosolic bacteriolysis.

Using a previously described luciferase-based assay, we confirmed that the Δ*menB* strain exhibited increased intracellular bacteriolysis (**Fig 5d**).^42^ We further found that NOX expression rescued Δ*menB* bacteriolysis, but not a comparable bacteriolysis phenotype in a Δ*glmR* strain that was previously shown to result from unrelated deficiencies in cell wall biosynthesis (**Fig 5d**).^52^ These studies thus show that efficient NAD^+^ regeneration is essential for limiting cytosolic bacteriolysis and suggest a model whereby respiration-mediated NAD^+^ regeneration promotes virulence, in part, by maintaining cell viability and facilitating evasion of innate immunity (**Fig 5e**).

## Discussion

Cellular respiration is one of the most fundamental aspects of bacterial metabolism and a validated antibiotic target. Despite its importance, the role of cellular respiration in systemic bacterial pathogenesis has remained largely unexplained. The studies reported here address the basis of respiration in the pathogen *L. monocytogenes*, identifying two electron transport chains that are partially functionally redundant and essential for pathogenesis. We find that restoring NAD^+^ regeneration to respiration-deficient *L. monocytogenes* strains through the heterologous expression of NOX prevents bacteriolysis within the host cytosol and rescues pathogenesis. These findings thus support the conclusion that NAD^+^ regeneration represents a primary role of *L. monocytogenes* respiration during pathogenesis.

Our results clarify several aspects of the basis and significance of energy metabolism in *L. monocytogenes*. In particular, our studies establish the relationship between *L. monocytogenes*’ two electron transport chains – confirming previous observations that flavin-based electron transfer enhances anaerobic *L. monocytogenes* growth and revealing a novel aerobic function of this pathway^25,27^. While the benefit of flavin-based electron transfer was only apparent in the absence of aerobic respiration, identifying the substrates and functions of aerobic activation of this pathway may provide an interesting avenue for future studies.

Our studies further reveal that *L. monocytogenes* employs a respiro-fermentative metabolic strategy that is characterized by production of the reduced fermentation products lactate and ethanol in the absence of an electron acceptor and acetate when a respiratory pathway is activated. This respiro-fermentative metabolism is consistent with the proton motive force being less central to *L. monocytogenes* energy metabolism and with a primary role of respiration in energy metabolism being to unleash ATP production via acetate kinase catalyzed substrate-level phosphorylation (**Fig 1f**).

The importance of cellular respiration for non-proton motive force-related processes is further supported by observations about the ability of heterologous NOX overexpression to rescue the severe pathogenesis phenotypes of respiration-deficient *L. monocytogenes* strains. NOX expression fully rescued *in vitro* growth defects and partially rescued virulence in the mouse model of disease, suggesting that NAD^+^ regeneration represents the sole function of respiration in some cell types and the major (but not sole) function of respiration in systemic disease. These findings suggest that a presently unaccounted for proton motive force-dependent aspect of microbial physiology is likely important for systemic disease. Considering the promise of cellular respiration as an antibiotic target, these insights into the role respiration plays in pathogenesis may inform future drug development strategies.

The centrality of NAD^+^ regeneration to *L. monocytogenes* also falls in line with relatively recent studies of mammalian respiration. Several studies have shown that the inability of respiration-deficient mammalian cells to regenerate NAD^+^ impacts anabolic metabolisms and inhibits growth^34,43–45^. Our discovery of a similar role of respiration in a bacterial pathogen thus suggests that the importance of respiration for NAD^+^ regeneration is a fundamental property conserved across the kingdoms of life.

## Methods

### Bacterial culture and strains

All strains of *L. monocytogenes* used in this study were derived from the wildtype 10403S (streptomycin-resistant) strain (see **Table 1** for references and additional details). The *Lactococcus lactis* water-forming *nox* (NCBI accession WP_010905313.1) was cloned into the pPL2 vector downstream of the constitutive P_hyper_ promoter and integrated into the *L. monocytogenes* genome via conjugation, as previously described.^46,47^ The Δ*qoxA*/Δ*cydAB*/*fmnB*::tn strain was generated from Δ*qoxA*/Δ*cydAB* and *fmnB*::tn strains using generalized transduction protocols with phage U153, as previously described.^48,49^

**Table 1.** Bacterial strains used in this study.

| Strains | Strain number | Reference |
| --- | --- | --- |
| *L. monocytogenes* (wildtype) | 10403S | 1 |
| $\Delta cydAB/\Delta qoxA$ | DP-L6624 | 2 |
| *fmnB::tn* | DP-L6612 | 3 |
| $\Delta cydAB/\Delta qoxA/fmnB::tn$ | DP-L7190 | This study |
| $\Delta fmnB::tn$ | DP-L7195 | This study |
| Wildtype + pPL2-NOX | DP-L7188 | This study |
| $\Delta cydAB/\Delta qoxA$ + pPL2-NOX | DP-L7189 | This study |
| $\Delta cydAB/\Delta qoxA/fmnB::tn$ + pPL2-NOX | DP-L7191 | This study |
| *Escherichia coli* | SM10 |  |
| NOX-pPL2 | DP-E7206 | This study |

1. Becavin, C. et al. Comparison of widely used *Listeria monocytogenes* strains EGD, 10403S, and EGD-e highlights genomic variations underlying differences in pathogenicity. MBio 5, e00969-00914, doi:10.1128/mBio.00969-14 (2014).
2. Chen, G. Y., McDougal, C. E., D'Antonio, M. A., Portman, J. L. & Sauer, J. D. A Genetic Screen Reveals that Synthesis of 1,4-Dihydroxy-2-Naphthoate (DHNA), but Not Full-Length Menaquinone, Is Required for *Listeria monocytogenes* Cytosolic Survival. MBio 8, doi:10.1128/mBio.00119-17 (2017).
3. Light SH, Su L, Rivera-Lugo R, Cornejo JA, Louie A, Iavarone AT, Ajo-Franklin CM, Portnoy DA. A flavin-based extracellular electron transfer mechanism in diverse Gram-positive bacteria. Nature. 2018 Oct;562(7725):140-144. doi: 10.1038/s41586-018-0498-z.

*L. monocytogenes* cells were grown at 37°C in filter-sterilized brain heart infusion (BHI) media. Growth curves were spectrophotometrically measured by optical density at a wavelength of 600 nm (OD_600_). An anaerobic chamber (Coy Laboratory Products) with an environment of 2% H_2_ balanced in N_2_ was used for anaerobic experiments. Media was supplemented with 50 mM ferric ammonium citrate or 50 mM fumarate for experiments that addressed the effect of electron acceptors on *L. monocytogenes* growth.

### Plaque assays

*L. monocytogenes* strains were grown overnight slanted at 30°C and were diluted in sterile phosphate-buffered saline (PBS). Six-well plates containing 1.2 x 10^6^ mouse L2 fibroblast cells per well were infected with the *L. monocytogenes* strains at a multiplicity of infection (MOI) of approximately 0.1. One hour post-infection, the L2 cells were washed with PBS and overlaid with Dulbecco’s Modified Eagle Medium (DMEM) containing 0.7% agarose and gentamicin (10 μg/mL) to kill extracellular bacteria, and then plates were incubated at 37°C with 5% CO_2_. 72 hours post-infection, L2 cells were overlaid with a staining mixture containing DMEM, 0.7% agarose, neutral red (Sigma), and gentamicin (10 μg/mL) and plaques were scanned and analyzed using ImageJ, as previously described.^49,50^

### Intracellular macrophage growth curves

*L. monocytogenes* strains were grown overnight slanted at 30°C and were diluted in sterile PBS. 3 x 10^6^ bone marrow-derived macrophages (BMMs) from C57BL/6 mice were seeded in 60 mm non-TC treated dishes containing 14 12 mm glass coverslips in each dish and infected with an MOI of 0.25 as previously described.^49,51^

### Mouse virulence experiments

*L. monocytogenes* strains were grown at 37°C with shaking at 200 r.p.m. to mid-logarithmic phase. Bacteria were collected and washed in PBS and resuspended at a concentration of 5 x 10^5^ colony-forming units (CFU) per 200 μL of sterile PBS. Eight-week-old female CD-1 mice (Charles River) were then injected with 1 x 10^5^ CFU via the tail vein. 48 hours post-infection, spleens and livers were collected, homogenized, and plated to determine the number of CFU per organ.

### NAD^+^/NADH assay

*L. monocytogenes* strains were grown at 37°C with shaking at 200 r.p.m. to mid-logarithmic phase. Cultures were centrifuged and then resuspended in PBS. Resuspended bacteria were then lysed by vortexing with 0.1-mm-diameter zirconia–silica beads for 10 minutes. Lysates were used to measure NAD^+^ and NADH levels using the NAD/NADH-Glo Assay (Promega, G9071) by following the manufacturer’s protocol.

### Fermentation product measurements

Organic acids and ethanol were measured by high-performance liquid chromatography (Agilent, 1260 Infinity), using a standard analytical system (Shimadzu, Kyoto, Japan) equipped with an Aminex Organic Acid Analysis column (Bio-Rad, HPX-87H 300 x 7.8 mm) heated at 60° C. The eluent was 5 mM of sulfuric acid, used at a flow rate of 0.6 mL/minute. We used a refractive index detector 1260 Infinity II RID and a 1260 Infinity II Variable Wavelength Detector (VWD). A five-point calibration curve based on peak area was generated and used to calculate concentrations in the unknown samples.

### Intracellular bacteriolysis assay

Bacteriolysis assays were performed as previously described.^30^ Briefly, immortalized *Ifnar*^-/-^ macrophages were plated at a concentration of 5 x 10^5^ cells per well in a 24-well plate. Cultures of *L. monocytogenes* strains were grown overnight slanted at 30°C and diluted to a final concentration of 5 x 10^8^ CFU per mL. Diluted cultures were then used to infect macrophages at an MOI of 10. At one hour post-infection, wells were aspirated and the media was replaced with media containing 50 μg/mL gentamicin. At six hours post-infection, media was aspirated and macrophages were lysed using TNT lysis buffer (20 mM Tris, 200 mM NaCl, 1% Triton [pH 8.0]). Lysate was then transferred to 96-well plates and assayed for luciferase activity by luminometer (Synergy HT; BioTek, Winooski, VT).

## Acknowledgments

Research reported in this publication was supported by funding from the National Institutes of Health (T32GM007215 to H.B.S., R01AI137070 to J-D.S., R01AI073843 & R01AI073843 to E.P.S, 1P01AI063302 & 1R01AI27655 to D.A.P., and K22AI144031 to S.H.L), the National Academies of Sciences, Engineering, and Medicine (Ford Foundation Fellowship to R.R.-L), the University of California Dissertation-Year Fellowship (to R.R.-L), and the Searle Scholars Program (to S.H.L). V.M.RR. holds a Postdoctoral Enrichment Program Award from the Burroughs Wellcome Fund and acknowledges support from the Academic Pathways Postdoctoral Fellowship at Vanderbilt University and the Howard Hughes Medical Institute Hanna H. Gray Fellows Program. Work at the Molecular Foundry was supported by the Office of Science, Office of Basic Energy Sciences, of the U.S. Department of Energy under Contract No. DE-AC02-05CH11231.

